# Tubulin as a sensitive target of nanosecond-scale intense electric field: quantitative insights from molecular dynamics simulations

**DOI:** 10.1101/533984

**Authors:** Paolo Marracino, Daniel Havelka, Jiří Průša, Micaela Liberti, Jack A. Tuszynski, Ahmed T. Ayoub, Francesca Apollonio, Michal Cifra

## Abstract

Intense pulsed electric fields are known to act at the cell membrane level and are already being exploited in biomedical and biotechnological applications. However, it is not clear if intra-cellular components such as cytoskeletal proteins could be directly influenced by electric pulses within biomedically-attainable parameters. If so, a molecular mechanism of action could be uncovered for therapeutic applications of such electric fields. To help clarify this question, we first identified that a tubulin heterodimer is a natural biological target for intense electric fields due to its exceptional electric properties and crucial roles played in cell division. Using molecular dynamics simulations, we then demonstrated that an intense - yet experimentally attainable - electric field of nanosecond duration can affect the *β*-tubulin’s C-terminus conformations and also influence local electrostatic properties at the GTPase as well as the binding sites of major tubulin drugs site. Our results suggest that intense nanosecond electric pulses could be used for physical modulation of microtubule dynamics. Since a nanosecond pulsed electric field can penetrate the tissues and cellular membranes due to its broadband spectrum, our results are also potentially significant for the development of novel therapeutic protocols.

**Author summary:** *α*/*β*-tubulin heterodimers are the basic building blocks of microtubules, that form diverse cellular structures responsible for essential cell functions such as cell division and intracellular transport. The ability of tubulin protein to adopt distinct conformations contributes to control the architecture of microtubule networks, microtubule-associated proteins, and motor proteins; moreover, it regulates microtubule growth, shrinkage, and the transitions between these states. Previous recent molecular dynamics simulations demonstrated that the interaction of the tubulin protein macrodipole with external electric field modifies orientation and conformations of key loops involved in lateral contacts: as a result, the stability of microtubules can be modulated by such fields. In this study, we seek to exploit these findings by investigating the possibility of fine-tuning the dipolar properties of binding sites of major drugs, by means of the action of electric fields. This may open the way to control tubulin-drug interactions using electric fields, thus modulating and altering the biological functions relative to the molecular vectors of microtubule assembly or disassembly. The major finding of our study reveals that intense (*>* 20 MV/m) ultra-short (30 ns) electric fields induce changes in the major residues of selected binding sites in a field strength-dependent manner.

## Introduction

Being able to control protein-based cellular functions with an electromagnetic field could open an exciting spectrum of possibilities for advancing biotechnological processes. In addition, it paves the way for the development of new biomedical theranostic approaches to treat various diseases where specific proteins are known targets. Electrostatic interactions in proteins are of paramount significance for protein function [1]. Strong molecular electric fields are known to play an essential role in protein folding [2], protein-ligand and protein-solvent interactions [3] as well as protein-protein interactions. Furthermore, the enzymatic activity of proteins [4] exploits local molecular electric fields to affect the potential energy surface of the reaction involved. A molecular electric field can also become crucial for protein stability since even a small imbalance in electrostatic interactions can cause malfunction of the protein [5]. Therefore, it is reasonable to assume that external electric fields (EFs) with appropriately chosen parameters of strength, frequency, and duration could modulate protein function. To overcome thermal noise effects and avoid heating side-effects, short (*<* 100 ns) intense (*>* MV/m) electric pulses lend themselves as a suitable form of electromagnetic field that can be utilized to modulate protein function [6, 7]. Indeed, it has been demonstrated through molecular dynamics simulations that EFs can affect conformation of pancreatic trypsin inhibitor [8], insulin [9–11], lysozyme [12–15], *β*-amyloid and amyloid forming peptides [16, 17], and soybean hydrophobic protein [18]. Further, EFs also unfolded myoglobin [19, 20], induced transition of peptides from a *β*-sheet to a helix-like conformation [21], and caused structural destabilization of (a short peptide) chignolin [22, 23] in molecular dynamics simulations. Moreover, recent studies demonstrated that EFs can affect water diffusivity and ion transport across transmembrane proteins such as aquaporins [24–28]. Experiments showed that EF applied directly by electrodes can catalyze the reaction [29] in a similar manner as a molecular electric field at enzyme sites [30] and switch protein conformational states [31]. Proteins from the tubulin family seem to possess an unusually high magnitude of charge and dipole electric moment [32, 33], thus possibly being a suitably sensitive target for the action of EFs. *α*- and *β*-tubulin monomers, which form stable heterodimers, are also crucial components of the cytoskeleton structures called microtubules, which are essential for cell division [34] among many other roles they play in living cells.

Several experimental works have been reported, which demonstrate effects of EFs on microtubule structures. It has been shown that EFs in the intermediate frequency range (100 – 300 kHz) had a profound inhibitory effect on the growth rate of a variety of human and rodent tumor cell lines [35] and also *in vivo* in the case of human brain tumors [36] presumably by interfering with the polymerization of mitotic spindle microtubules which are composed of tubulin dimers. In a recent work, an ultra-short intense pulses EF cause Ca^2+^ independent disruption of dynamic microtubules in glioblastoma [37]. However, it is not clear whether the observed effect of EFs was direct or indirect through the action on the membrane channels first and then transmitted downstream into the cell interior. Hence, an exact molecular-level mechanism of this electric field action on microtubules and tubulin remains unknown.

Microtubules, due to their expected special electric [38] and vibrational [39–42] properties, were proposed to be involved in endogenous electrodynamic processes in cells [43–45]. However, all-atom molecular simulations of external EF effect on tubulin have been carried out only recently [46, 47]. They specifically investigated the EF effects on protein mechanics but did not include the C-terminal tail. The C-terminal tail is a highly flexible unstructured domain of tubulin, which (i) is essential for tubulin-protein interactions [48, 49], (ii) is the main site of protein mutation and post-translational modifications [49], and (iii) greatly contributes to the overall electric properties of tubulin accounting for approximately 30-40% of the total charge [33]. A very recent study [50] presented results of the molecular dynamics simulation analysis of the electric field effect on tubulin. This latter study investigated electric field strengths in the range between 50 and 750 MV/m, which overlaps with the values used in our simulations, and carried out simulations over 10 ns while our study reports on simulations that ran up to 30 ns. The previous study only examined displacement effects on key secondary structure motifs such as tubulin’s C-termini and alpha helices. In contrast, in the present study, we aim to unravel the mechanisms of interaction of intense EFs on the tubulin protein family, given their important and attractive role as drug targets for cancer therapy, due to their involvement as key-players in cellular self-organization. In fact, it is already known that various stabilizing/destabilizing tubulin-binding drugs such as taxanes, colchicines, and vinca alkaloids, bind to different sites on the tubulin dimer, modulating microtubule-based processes [51]. This type of binding is assumed to be based on purely electrostatic interactions like those employed in the recognition between proteins [2]. Conversely, what is presently unexplored, is the possibility to modulate such electrostatic environment by means of EF. Therefore, we first employ bioinformatics tools to systematically compare electric charges and dipole moments of the various isoforms of tubulin to the so-called PISCES set, which represents all unique chains in the whole protein database. Then, we quantitatively evaluate the effects of intense nanosecond EFs on the overall shape of the tubulin dimer as well as the time-dependent evolution of its dipole moment. Finally, we closely analyze the dipole moments of individual residues with special attention to those forming the binding pockets for the most important tubulin-modulating pharmacological agents (e.g. paclitaxel or vinca alkaloids) and for GTP/GDP molecules. Thus, in this paper, we cast light on the mechanism of direct EF action on tubulin at the molecular level.

## Results

### Proteins from tubulin family have exceptional electric properties

In this study we first compared the structural charges and dipole moments of proteins from the tubulin family to those of the PISCES set of proteins (all unique protein chains available in the RSCB protein database). It can be seen from Fig. 1A that tubulin proteins possess a generally much higher structural electric charge (with a mean value of −22 e per monomer) than most PISCES proteins (mean value of −4 e). This is likely due to an excess of acidic over basic amino acid residues present among the tubulin proteins compared to the PISCES set of proteins, see S1-1TUB-aa content.xlsx. Taking as an example the 1TUB tubulin structure, we found that while a relative content of acidic residues is similar in tubulin versus the average from the PISCES set, tubulin contains fewer basic residues than an average protein. It is worth noting that a large fraction (on average around 34% across the tubulin dataset) of electric charge of tubulin is due to the unstructured and highly flexible C-terminal tail (CTT). This is remarkable, since the CTT of tubulins is rather short - having on average around 20 residues. A rather high value of the electric charge of tubulin CTT is due to a large content of acidic residues.

**Fig 1.**
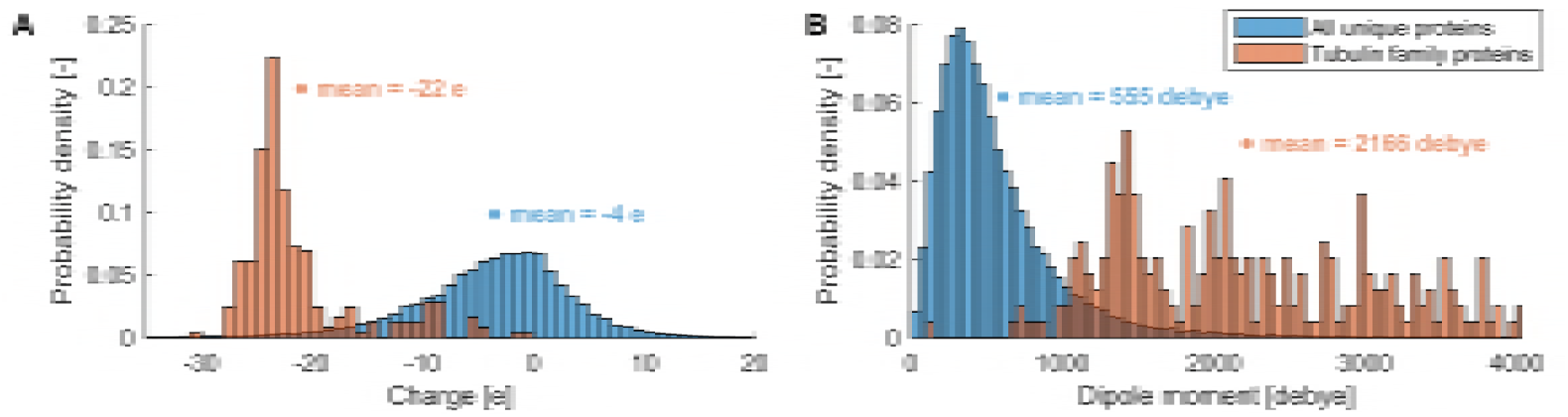
Structure of proteins from the tubulin family indicates that they have exceptional electrostatic properties compared to other proteins. (A) distribution of electric charge values for the tubulin family (orange) and the PISCES set of approximately 63,000 unique protein chains (blue). (B) distribution of dipole moment values for the tubulin family (red) and the PISCES set of approximately 63,000 unique proteins chains (blue). *•* depicts a horizontal position of a mean value.

Fig. 1B shows that tubulins possess a much higher dipole moment than PISCES proteins: with a mean of (2,166 debye) versus a mean of (555 debye), respectively. The high dipole moment arises from an asymmetric electric charge distribution, i.e. an asymmetric distribution of acidic versus basic residues. This asymmetric charge distribution is partially due to the fact that a large fraction of charge is located on the C-terminal tail. However, even the tubulin structures where the CTT is not resolved possess a rather high dipole moment, e.g. both 1TUB and 1JFF crystal structures result in a dipole moment above 2,000 debye. The results in Fig. 1 clearly demonstrate that the tubulin proteins have exceptional electric properties, hence it is reasonable to analyze in detail if these properties can be exploited to manipulate tubulin’s structure and hence its function using an external EF with specifically designed characteristics.

### Rounding effect on tubulin shape induced by electric field

We have employed molecular dynamics (MD) simulations, see Methods for details, to analyze effects of an EF acting on tubulin. The tubulin structure used in these simulations consists of *α*- and *β*-monomers, which form a stable noncovalently bonded heterodimer, see Methods for details.

In the MD simulation performed, we have added an EF as an external Coulomb force acting on every atom in the system. We analyzed the effect of the EF strength starting from 20 MV/m up to 300 MV/m. The rationale for the selection of this range of field strength is both theoretical and experimental. Theoretically, the interaction energy *U* = ***p · E***, where ***p*** is the tubulin dipole moment vector and ***E*** the electric field vector, exceeds thermal energy for field strengths in the range *>* MV/m. The range *<* 70 MV/m is also experimentally attainable since it is below the field strength of dielectric breakdown of water-like media with an exact value depending on the ionic strength and the electric pulse duration [52]. The length of the simulation was selected to be up to 30 ns. This time scale is short enough to be computationally tractable and long enough to be comparable to experimental data. Furthermore, electric pulses of this duration typically do not cause any appreciable heating in experiments conducted even for the experimentally attainable field strengths mentioned above.

The most basic integral parameter of protein is its shape. Hence, we first analyzed how the EF affects the shape of the tubulin dimer, see Fig. 2 (note that no water molecules and counterions are displayed). To this end, we approximated the shape of tubulin by an ellipsoid and obtained a new coordinate system within the frame of this ellipsoid, see Methods. This approach allows a sound analysis of protein polarization process, bypassing the trivial roto-translational effects taking place due to the external EF. At first, we observed that the effective shape of the equivalent tubulin ellipsoid is affected by a 100 MV/m EF in a way that the medium axis is elongated by 11% and the long axis is shortened by 2%. We also observed that tubulin undergoes rotation - this will be discussed in more detail in the next section as the rotation is related to the dipole properties of the tubulin analyzed there.

**Fig 2.**
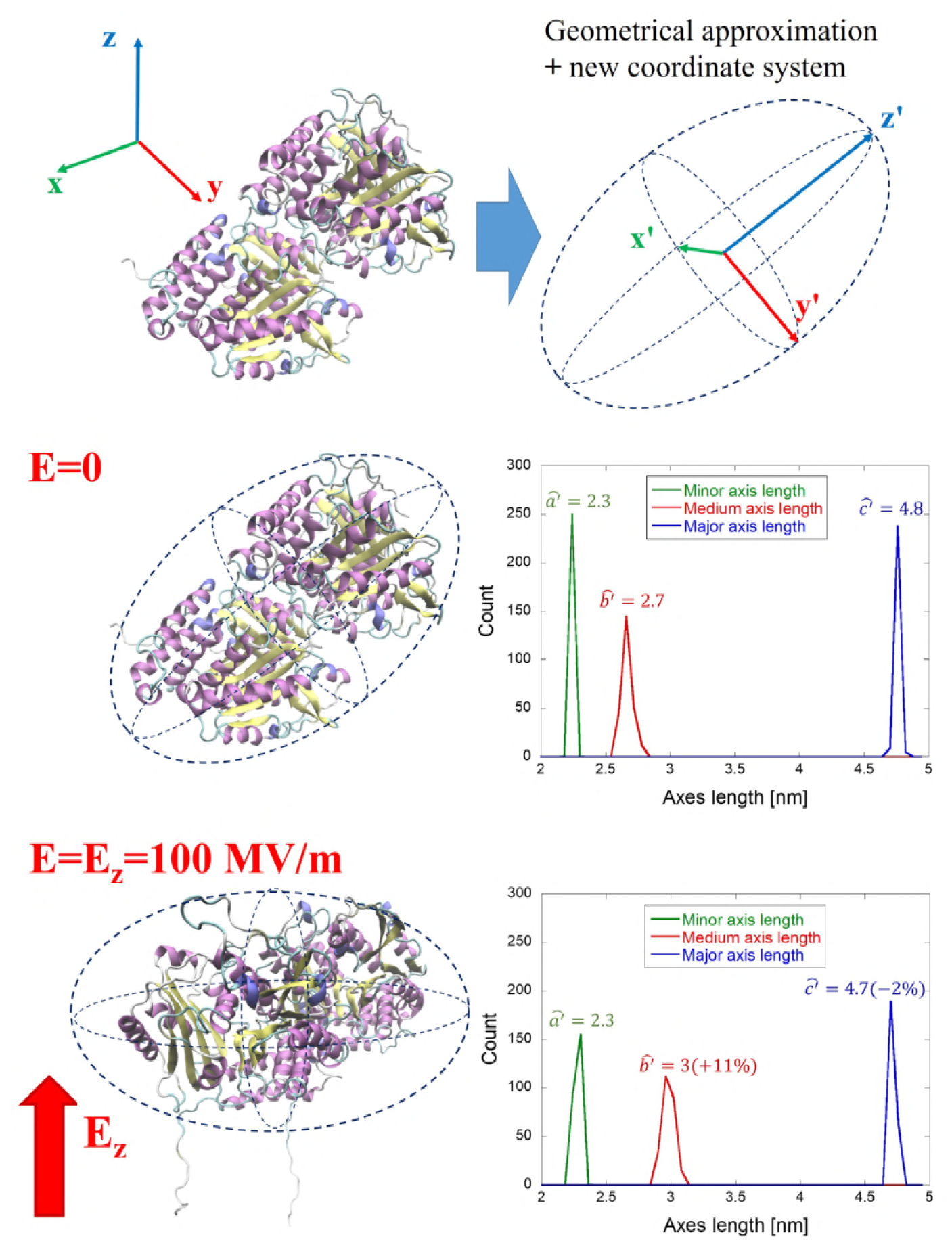
New coordinate system and dimensions of the tubulin approximated by an ellipsoid. The electric field affects not only the orientation of the tubulin dimer but also its overall shape. The distributions on the right are from 2,500 frames from the last 5 ns of the MD simulation (sampling rate 2 ps).

### Orientation effect of electric field on tubulin dipole moment

As the next step in this investigation, we focused on the effect on the electric dipole moment of the tubulin dimer, see Fig. 3. It is readily seen that the tubulin dipole moment under zero-field conditions has an average value of around 2,500 debye. However, in the presence of a 100 MV/m EF, the dipole moment is increased to more than 6,500 debye, i.e. it more than doubles in the process. In the lower part of the figure it is also visible the polarized C-terminal tails extending from the structure. To get deeper insight into time evolution and field strength dependence, we further analyzed the tubulin y′ dipole component, the most affected one (i.e. with the highest polarization change) among the three dipole components shown in Fig. 3. The external electric field vector was oriented in the z direction (Cartesian reference system, see Fig. 2 lower-left panel) during the whole course of the MD simulation while the initial orientation of the tubulin dipole had an anti-parallel component to it. At the zero electric field strength the effective y′ component of the tubulin dipole moment fluctuates around 2,000 debye, with a rather steady profile for the whole simulation time (see Fig. 4). This stable dipole value evolution is due to the coordinate system transformation adopted (i.e. from the external Cartesian system to the internal protein system), which removes the dipole roto-translational effects, highlighting internal polarization effects. In fact, the EF tends to act by exerting a torque on the tubulin’s dipole moment so that the dipole component in the direction of the field vector is maximized. Data reported in Fig. 4 suggest that EF in the range of 20 MV/m induce a slow, but significant, polarization in the last 5 ns of simulation, so that the y′ dipole component increases from 2,000 to 3,000 debye. Higher external fields amplify and speed up this polarization effect up to a value above 5,000 debye after 7 ns for a field strength of 100 MV/m. Moreover, a second field-effect appears on the *β*-tubulin CTTs. While in the zero field simulation and within approximately 20 ns of 20 MV/m condition, the CTT remains more or less close to the surface of tubulin, for the field strengths ≥ 50 MV/m the CTT is pulled away from the tubulin surface and becomes outstretched (see Fig. 3).

**Fig 3.**
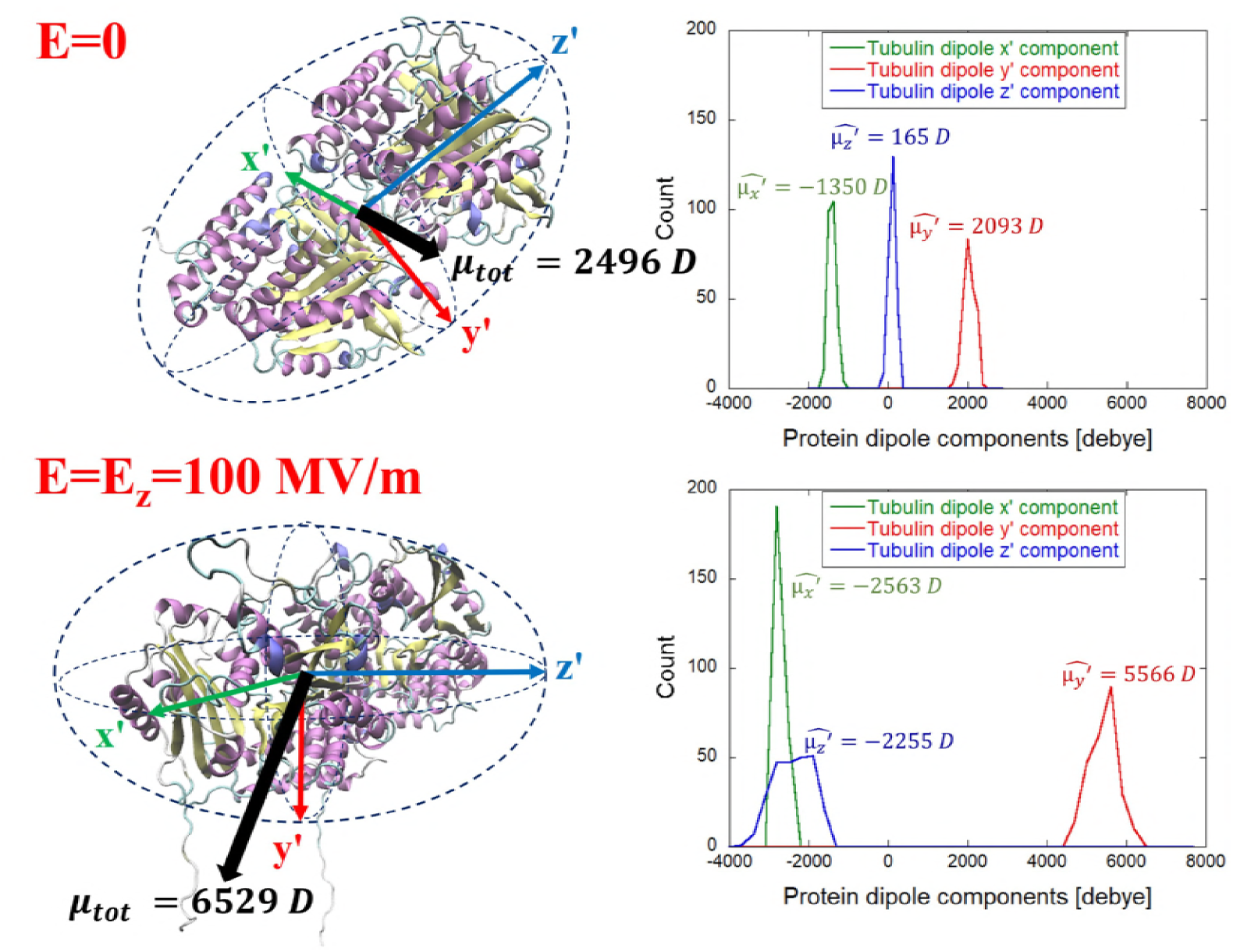
Electric field affects not only tubulin orientation but also the overall tubulin dipole moment. The distributions on the right are from 2,500 frames from the last 5 ns of the MD simulation.

**Fig 4.**
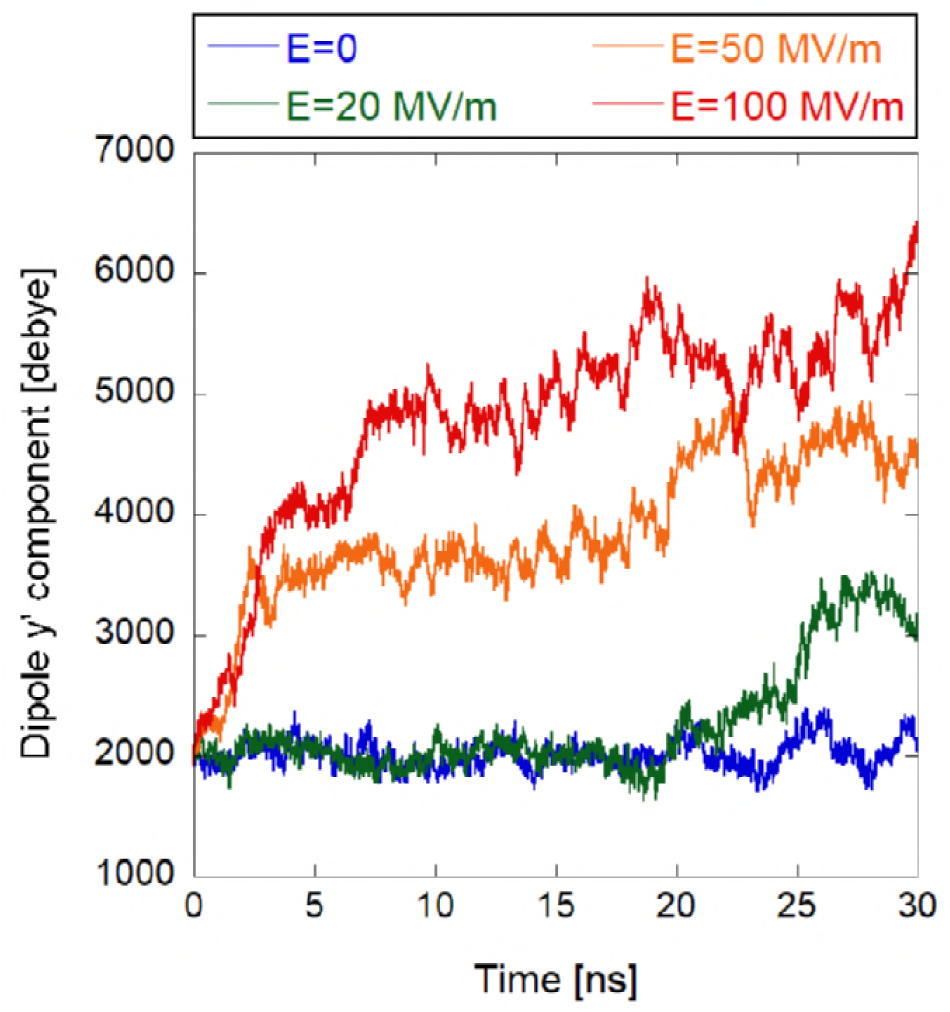
Evolution of the y′ component of the tubulin dipole moment.

### No unfolding effect on tubulin up to 100 MV/m

The fundamental and functionally-crucial characteristics of any protein structure are the type and location of its secondary structure motifs. Therefore, we have focused next on the analysis of how an EF affects the count of tubulin residues being part of a certain secondary structure motif. The results of this analysis can be seen in Fig. 5, which displays the count of residues in alpha helices, beta-sheets, coils and turns as averaged over the last 5 ns of the corresponding trajectories. The field strength of 100 MV/m seems not to cause any effect on the structure - just slightly increasing the content of the tubulin residues present in coils at the expense of beta-sheets, alpha helices and turns. Only 300 MV/m induces a strong effect on tubulin secondary structures, where more than 70 residues are converted from alpha-helices to coils. This is a sign of unfolding, when charged CTTs are experiencing an electrostatic force, which is strong enough to undergo pulling out from the tubulin body. Since there are two alpha helices (H12 and H11) close to CTTs, those are the first ones which are unfolded. Furthermore, we selected the CTT of *β*-tubulin since it is longer and carries more charge than the *α*-tubulin CTT for a more detailed view. In Fig. 6A, we quantified the count of CTT residues being in coil structures within the 30 ns time scale for 0, 20, 50, and 100 MV/m field strength conditions. We can see that the 20 MV/m condition does not tend to change the secondary structure of CTT compared to the no field condition. However, both 50 and 100 MV/m field strengths tend to increase the count of residues being in a coil structure at the expense of *α*-helices and turns. In these cases, the higher the field strength, the shorter the time needed to achieve the maximum amount of residues in a coil structure.

**Fig 5.**
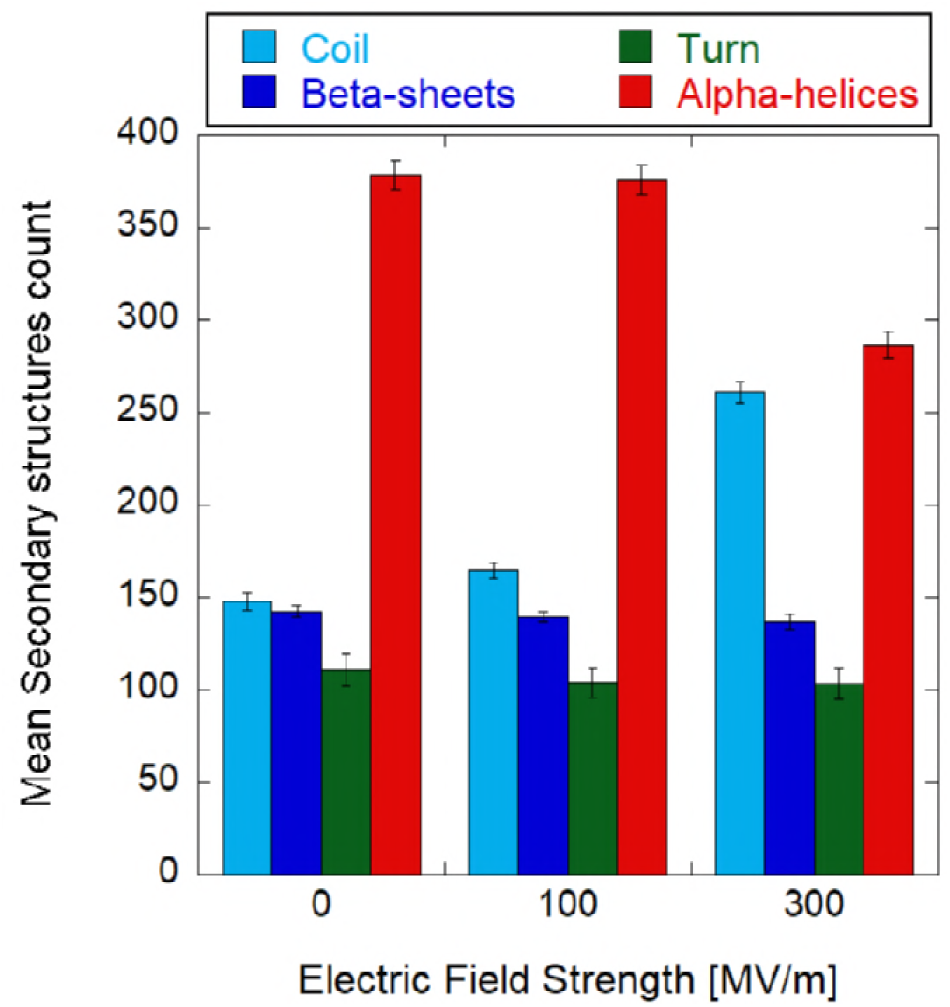
Tubulin’s main secondary structures under three different exposure conditions. Error bars represent the standard deviations as obtained by the last 5 ns of the corresponding equilibrium trajectories.

**Fig 6.**
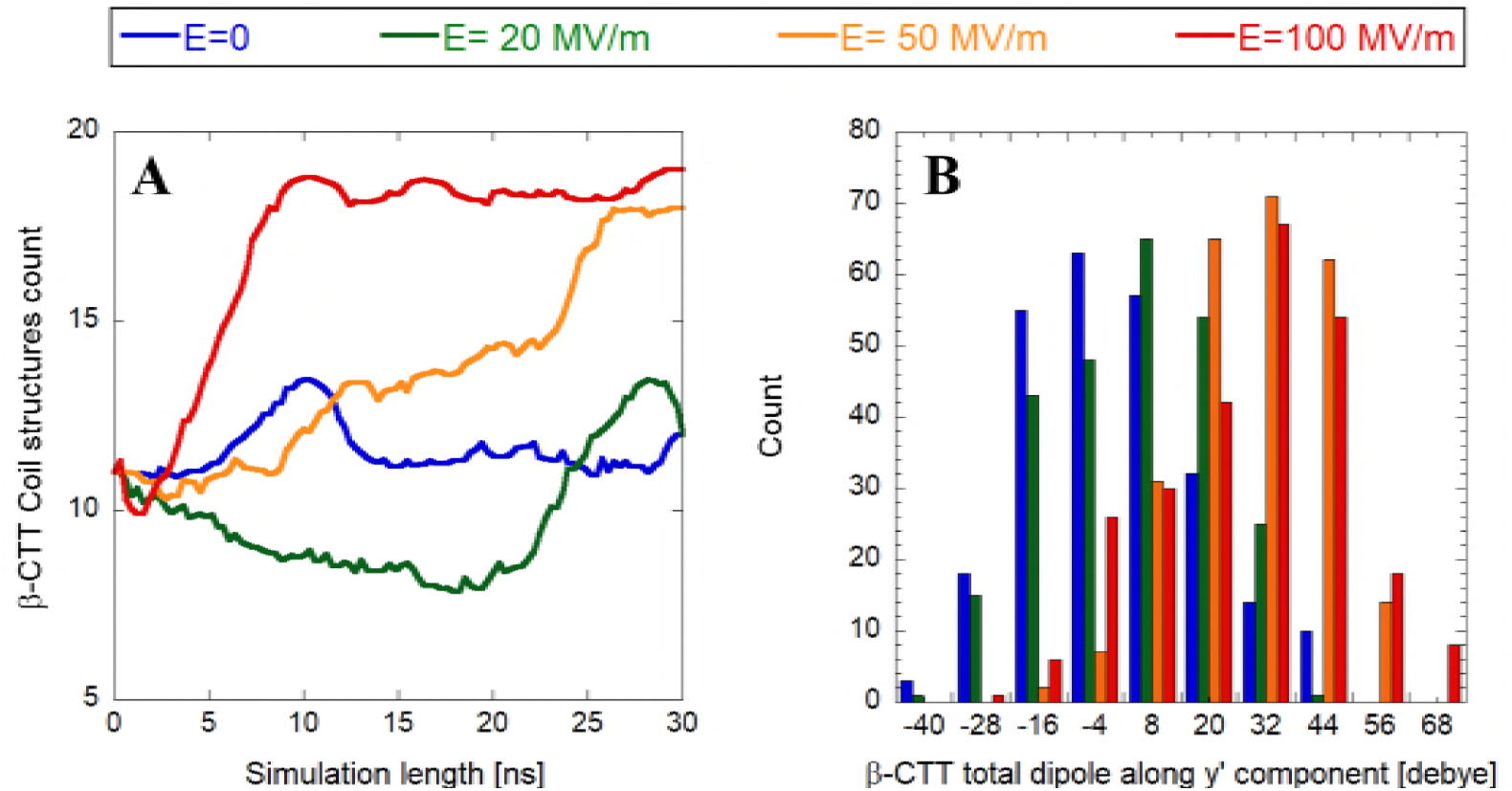
Panel A: smoothed representation of *β*-tubulin tail coil secondary structures; panel B: histogram representation of electric field effects (ranging from zero up to 100 MV/m) on the whole *β*-tubulin tail dipole component along the y′ internal coordinate (data refer to the last 5 ns of each simulation).

### Electric field affects the dipole moment of specific tubulin domains

C-terminal tails of tubulin are important for interactions with microtubule-associated proteins, such as motor proteins [53]. Since the data in Fig. 3 suggested substantial effects on CTTs of tubulin, we next focused on a detailed analysis of CTTs. In panel B of Fig. 6, we found a strong polarization effect on the CTT of *β*-tubulin, with a mean dipolar shift for field strength ≥ 50 MV/m as high as 40 debye.

Apart from the EF effect on CTTs, we also asked a question whether an intense field can affect local electrostatic conditions of the tubulin dimer. To this end, we analyzed the shift of the dipole moment of every residue of the tubulin dimer. We plot the y′ component (internal tubulin coordinates) of the dipole moment in Fig. 7. It can be seen that some residues undergo substantial change of the dipole moment when exposed to an EF of 100 MV/m. In particular, when considering both monomers it is possible to appreciate dipolar shifts ranging from −10 up to almost +15 debye. Specifically, 22 out of the 451 residues of *α*-monomer residues exceed a ±5 debye shift, while for the *β*-monomer the concerned residues are 3 out of 445 ones. To understand if dipole moments in any functionally significant parts of the tubulin dimer are affected, we focused on selected important sites of the tubulin dimer: GTP binding/hydrolysis site, sites of tubulin longitudinal interactions, and the binding sites of the three most common tubulin drug families (paclitaxel, nocodazole/colchicine, and vinca alkaloids). The Methods section provides a rationale and identification of residues belonging to the tubulin sites based on energy considerations. In Fig. 8, we highlight the location of the important tubulin sites in a color-coded manner and also provide a list of the selected residues. We display the histograms of the dipole moments (y′ component) of six selected residues where we found the strongest effects. In all of these six cases, we see a field strength-dependent effect, shifting the y′ component of a residue dipole moment towards different values. Since the local electrostatic field is crucial in active sites of many enzymes enabling the process of catalysis, we speculate that influencing the local field could affect the GTP hydrolysis rate. We see that *α*-Asp 251, a crucial residue mediating GTP hydrolysis [54], has its y′ component of the dipole moment (yDM in short) affected by almost 3 debye when comparing no field and 100 MV/m conditions. Tubulin-tubulin longitudinal interactions are mediated by several residues. *β*-Arg 391 is one of them and has its yDM also influenced by the EF (Fig. 8). However, in this case the field strength has opposite sign effect. 20 MV/m field tends to affect yDM in an opposite direction than fields with 50 and 100 MV/m field strengths, compared to no field condition. This is probably due to the position of the residue in the *β*-monomer (see Fig. 8, left-upper panel); in fact Arg-391 is located in the external part of the protein, presumably in contact with a layer of water, therefore its response to the external EF could be screened by water for lower field strength. The second most important residue for binding paclitaxel energy-wise, *β*-Val 23 (Fig. 8), also manifests the changed yDM up to few debye when an EF of 100 MV/m is applied. Furthermore, residue *β*-Cys 239 belonging to the colchicine binding site has its yDM shifted towards zero with an increasing field strength. We also found that *α*-Pro 325 and *β*-Asp 177 belonging to the binding site of the vinca alkaloids family are influenced by an EF. In this case however, the shift of yDM is only in the sub debye range.

**Fig 7.**
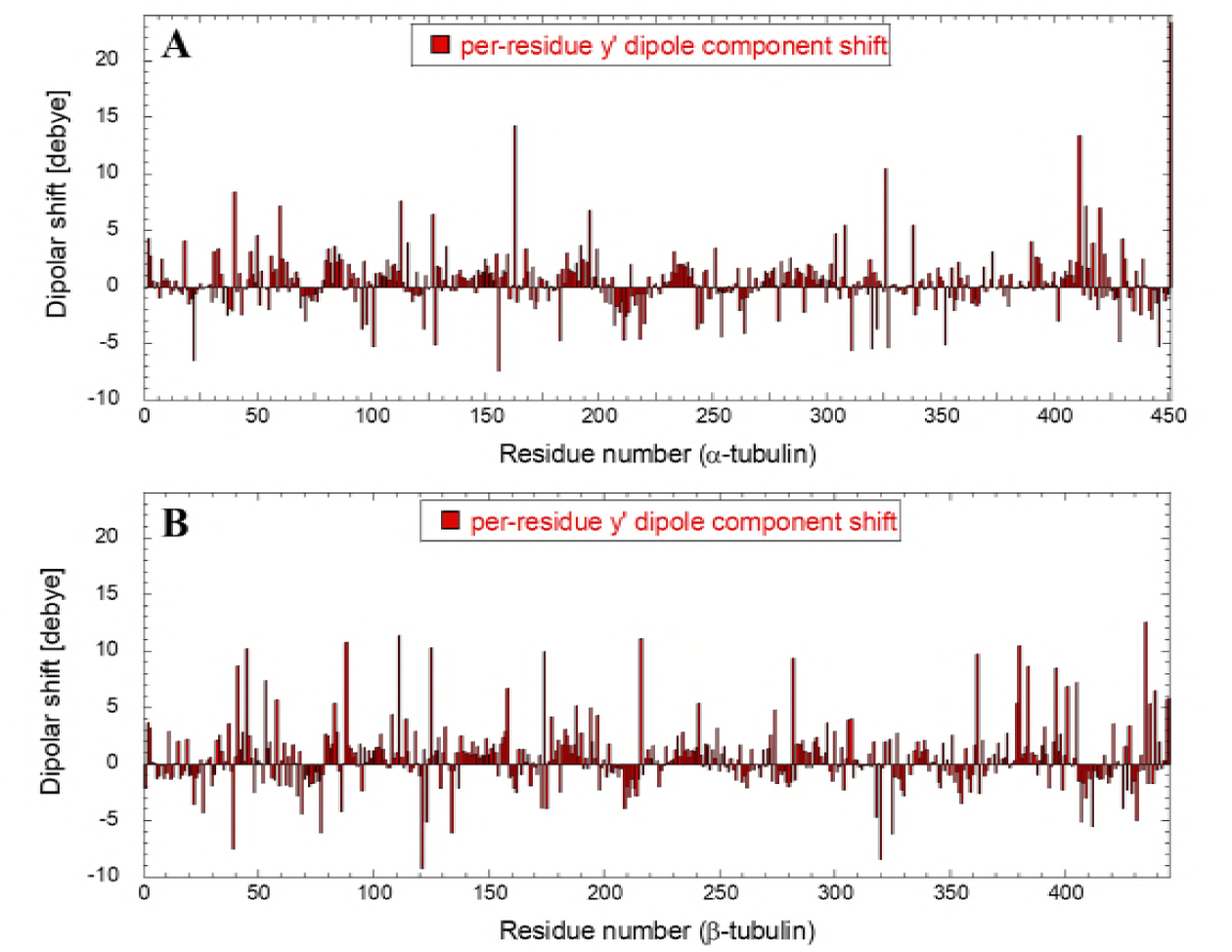
Dipolar shift of each tubulin residue (Monomer A in panel A, monomer B in panel B) in the presence of the 100 MV/m electric field strength with respect to the unexposed case. Data refer to the dipole component along the y′ internal coordinate (see S2-x,z dipole components for the complete picture of dipole component shifts).

**Fig 8.**
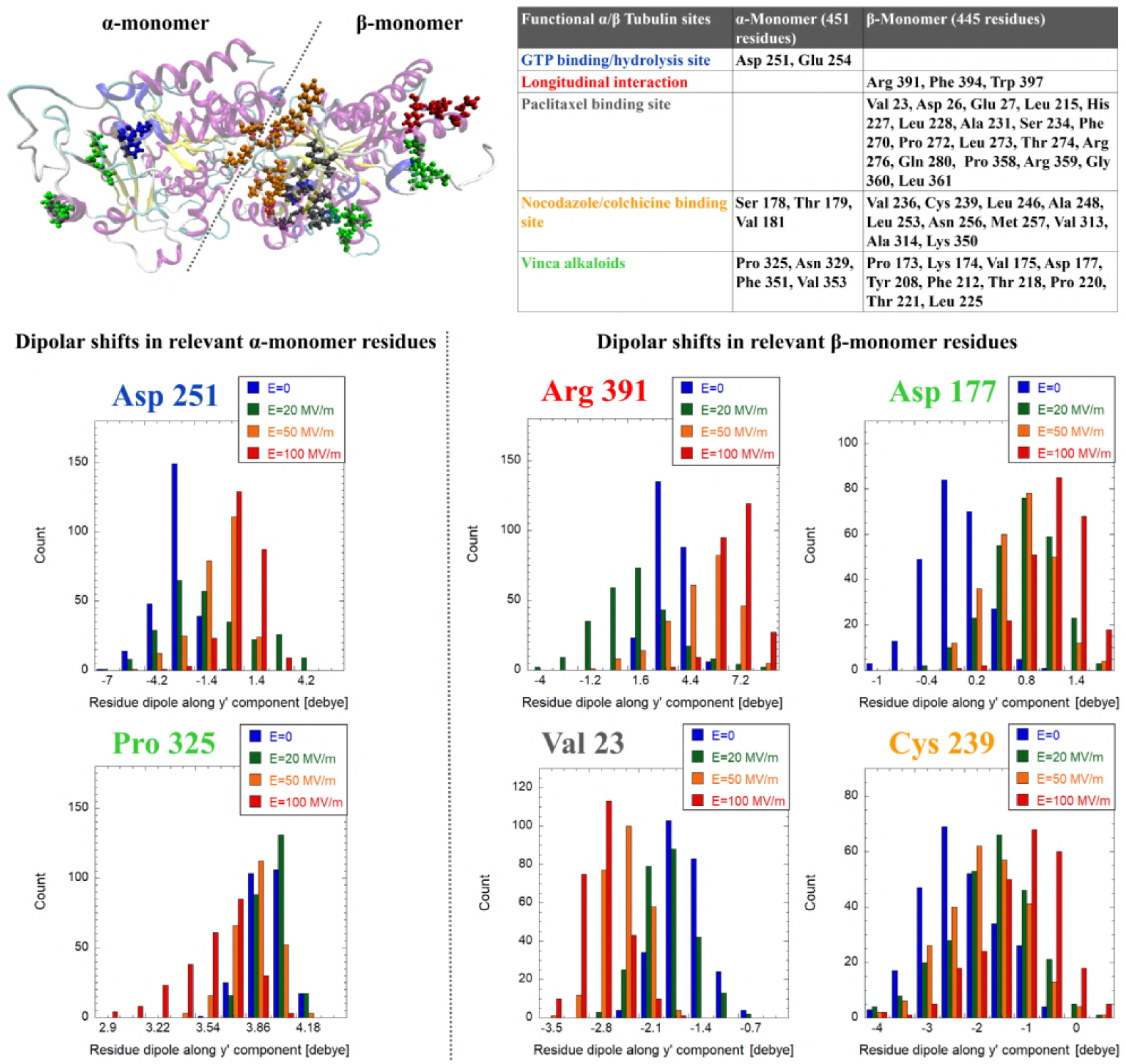
Electric field effect on selected important tubulin residues; in the upper panel a graphical representation of known *αβ*-tubulin sites is given together with a look-up table - see text criteria and for references where the residue numbers were obtained. In the lower panel, the histogram representation of electric field effects (ranging from zero up to 100 MV/m) on a dipole moment of selected relevant residues is presented calculated from the frames of the last 5 ns of simulation.

## Discussion

We have demonstrated that by using MD simulations combined with the Covariance Matrix method, it is possible to study with high precision the intrinsic structural and dipolar response of tubulin protein under the action of nanosecond scale EFs. The findings presented here cover several aspects of such tubulin/EFs interaction mechanisms. First, we quantified the unusually high magnitude of structural charge and dipole electric moment of tubulin family proteins in absence of EFs and, specifically, the response of tubulin to nanosecond scale EFs. Second, we studied structural/conformation changes and possible unfolding effect on the whole protein. Finally, we provided evidences of dipolar coupling with the EF of residues forming the binding pockets for the most important tubulin-modulating pharmacological agents (taxane, vinblastine, and colchicine) as well as tubulin CTTs. In the following we discuss the significance of such results, as well as the limitations and point out some possible extensions for future work.

### Electric and dipolar properties of tubulin

Regarding the charge and dipolar properties of tubulin family, we shown that an average (across the list from various species) tubulin monomer has an approximately 4-5 fold higher structural electric charge and electric dipole moment than an average protein (Fig. 1). This result corroborates our further findings: tubulin seems to be more susceptible to EFs compared to other proteins analyzed earlier [11, 12, 19], in a way that lower field strength, i.e. 20 MV/m, is able to attain a 50% increase of tubulin dipole.

In a recent paper [55] a detailed energy balance calculation was provided for the stability of a microtubule with a seam (representing a so-called type B microtubule lattice). The calculations included the contributions from dipole-dipole interactions between tubulin dimers, solvent accessible surface area, van der Waals and electrostatics (generalized Born approximation) to demonstrate that the balance energy is such that a single GTP hydrolysis event can trigger a microtubule disassembly because when the seam is closed with GTP molecules attached to the *β* monomers, the net free energy is -9 kcal/mol. The dipole-dipole energy is positive (destabilizing) and amounts to 27 kcal/mol. When the seam becomes open due to GTP hydrolysis, the net free energy becomes vastly positive. These calculations used a dipole moment of approximately 4,500 debye for a tubulin dimer as an average value over various isotypes. In the present paper we have shown that strong electric fields can substantially increase the dipole moment of tubulin. Here, we have shown the dipole moment to be close to 3,000 debye in zero field, which would translate into a dipole-dipole energy for an microtubule lattice of 12 kcal/mol and a net free energy of −24 kcal/mol, hence slightly more stable than the calculation provided in [55]. However, at strong electric field values (≥ 50 MV/m), the corresponding dipole moments were found to increase to as much as 6,000 debye, which translates into a dipole-dipole energy of 48 kcal/mol and a net free energy of +12 kcal/mol making the lattice unstable. This quantitatively supports our hypothesis that sufficiently strong electric fields disrupt a microtubule lattice by increasing the dipole moments of tubulin and contributing a positive energy that cannot be balanced by the remaining contributions.

### Structural/conformation changes on tubulin induced by electric fields

By approximating the tubulin with the ellipsoid as given by the covariance Matrix method, we were able to appreciate the actual shape changes under the action of the EFs. The protein slightly reduces its major axis, while clearly increasing its medium one. This effect seems of particular interest since it is a consequence of protein polarization. In fact, the tubulin dimer undergoes a packing transition, rather than an elongation, since its dipole moment aligns along tubulin medium axis (see Fig. 3). This non-trivial effect is due to the high charge of tubulin CTTs. Tubulin seems to have a lower threshold for the unfolding transition [56]. For example, unfolding effects tend to appear at lower field strengths or within a shorter time frame (unfolding at 250 MV/m started at approximately 160 ns in insulin [11, Fig.1c] compared to only a few ns at 200 MV/m for tubulin). This is reasonable since the higher the charge and dipole moment of the structure, the higher the electric force and torque, respectively, acting on the protein.

### Dipolar coupling of specific tubulin domains with the electric field

We found that a primary target of electric field in tubulin heterodimer are C-terminal tails since they carry a substantial amount of electric charge. Indeed, they manifest a strong polarization effect on the CTT of *β*-tubulin, with a mean dipolar shift for field strength ≥ 50 MV/m as high as 40 debye. We might ask a question: how biologically general is the effect on the CTT we observe, or put another way, is the CTT sequence we used common in other biological species?

An exact tubulin sequence and structure varies across biological species and tissues [33]. While the tubulin core is rather conserved across species, CTTs display higher variability across species and are also a target of post-translational modifications forming a so-called “tubulin code” [57, 58]. The *β*-tubulin CTT sequence we used in our simulations has a 100% sequence identity to certain tubulin isotypes found for example in pig *Sus scrofa* (common experimental source of tubulin) tubulin gene TBB and to many tubulin isotypes of more than 20 species, see S3-CTT alignment.fasta. It has also a 90% sequence similarity to CTT of TBB2B gene in the human, the only difference is that two immediate neighbor residues Gly 440 and Glu 441 are swapped, so the total charge of the CTT is the same as that of the CTT in our tubulin structure, see S3-CTT alignment.fasta.

As a general result, we observed a widespread EF effect on protein residues. In particular, when analyzing specific tubulin domains, e.g. the GTP binding/hydrolysis site, sites of tubulin longitudinal interactions, and the binding sites of the three most common tubulin-binding drug classes (paclitaxel, nocodazole/colchicine, and vinca alkaloids), we obtained unexpected dipolar coupling even in the presence of a 20 MV/m acting for 30 ns. In this context it is interesting to note that at present, generators able to produce intense (*>* 20 MV/m) ultra-short (30 ns) electric field pulses, are commercially available, hence the intriguing perspective to control and modulate protein functions is becoming realistic. For instance, one can envisage that pulsed electric field could be used in synergy with taxane-based cancer treatment protocols to modulate the drug binding to tubulin and hence the dynamics of microtubules. It is worth noting that the most negatively charged part of the tubulin dimer, when embedded in a microtubule, is the cytosol-facing outer surface while the surface facing the lumen is less negatively charged. Also, the dipole moment of the dimer also includes a contribution from the CTT which is highly variable due to the flexibility of this motif. Finally, the dielectric breakdown, which would pose a limitation on the strength of the applied field in practice, depends on the ion strength of the solution and the duration of the applied pulse [52].

### Limitations

In discussing the implications of our modeling effort for experiments on microtubules, it could be argued that field strengths *>* 100 MV/m are not readily experimentally attainable yet. However, one has to keep in mind that MD simulations are commonly best suitable for providing relative assessment of various effects and not necessarily absolute values directly translatable the experimental situation. For example, the binding free energies of ligands interacting with proteins are typically off by a factor as large as two or more [59]. Hence, effects such as unfolding, which are computationally predicted to occur at large fields, may in fact require field strengths lower than predicted for tubulin [56]. Moreover, rapid technological advances in the field may well lead to the development of engineering solutions with sufficiently high electric field strengths attainable in the clinical setting. The ultimate physical limit, however, is the field strength at which the dielectric breakdown of the exposed biological material occurs.

### Challenges for future work

In our future work we intend to better calibrate the simulation parameters in order to make the model quantitatively predictable so that such characteristics as field strength, frequency and duration may be tailor-designed to elicit specific response of the target, namely the tubulin dimer or an entire microtubule. This modeling effort may need to be extended to include quantum effects through the use of quantum mechanics/molecular mechanics calculations especially to be able to elucidate covalent bond response to electric fields, for example at the GTP binding site.

To conclude, our results allow to draw biologically-relevant consequences in terms of microtubule stability and the changes in the strength of binding for tubulin-binding drugs under the influence of EF. Therefore, we believe the present paper provides new and important quantitative insights into the effects of EF pulses of nanosecond duration on tubulin and microtubules. The knowledge of residue-specific EF realignments allows to extrapolate the computer simulation results in terms of their consequences on the behavior of other tubulin isoforms and tubulin mutants under both exogenous and intrinsic molecular electric field.

## Methods

We follow a gapless numbering convention of tubulin residues in the current paper, in contrast to the Protein Data Bank entries 1JFF and 1TUB, which contain 2 gaps after *β*-Leu 44 and 8 gaps after *β*-Pro 360.

### Comparison of tubulin family proteins to PISCES set of proteins

Charge and dipole moment of proteins from tubulin family was obtained from our earlier work [33], which includes 246 isotypes of tubulin, mostly *α*- and *β*-tubulin, from various species. PISCES set, a representative set of all proteins in the RSCB PDB database, was obtained using PISCES server [60]. We used following settings to run the server query (date 2018-05-18): sequence percentage identity threshold ≤ *90*, Resolution: *0.0 – 4.0*, R-factor: *0.5*, sequence length: *40 – 10,000*, Non X-ray entries: *Included*, CA-only entries: *Excluded*, Cull PDB *by entry*, Cull chains within entries: *No*. We obtained a list with 36,331 entries, see S4-PDB entries.zip. Charges and dipole moment data for our PISCES set of proteins were obtained from Protein Dipole Moments Server of Weizmann Institute https://dipole.weizmann.ac.il/index.html [61]. Each protein chain was evaluated separately which leads to a total number of 63,110 records for both protein chain charge and dipole moment, see S5-PISCES-Charge, Dipole.zip. The effective pH considered is 7 [61].

Note that the methods for charge and dipole moment calculation used by Tuszynski et al. [33] (GROMOS96 43a1 force field) and the dipole moment server [61] are not the same. However, when we tested both methods on selected tubulin structures 1TUB and 1JFF, they yielded very similar values: 1TUB charge −34 vs. −33 e, dipole moment 1,998 vs. 2,040 debye; 1JFF −31 vs. −31 e, dipole moment 1,645 vs. 1,666 debye; for the method by Protein Dipole Moments Server vs. Tuszynski et al. [33].

### Tubulin structure

The structure of the GMPCPP-bound tubulin was obtained from the Protein Data Bank under the PDB ID: 3J6E [62]. The cofactor GMPCPP, a non-hydrolyzable analogue of GTP, was modified into GTP. Furthermore, the CTT, which are usually missing in PDB structures of tubulin, were added to both *α*- and *β*-tubulin subunits. This was achieved using the Molecular Operating Environment software (MOE, Chemical Computing Group Inc.) [63]. MOE was also used for the addition of hydrogen atoms and the prediction of ionization states. The CTTs were added in an extended conformation and we depended on molecular dynamics simulations later to help to restructure the tails in the correct conformation in solution. The CTT is defined here as the last 16 residues of the *α*-tubulin (GVDSVEGEGEEEGEEY) and the last 20 residues of the *β*-tubulin (QDATADEQGEFEEEGEEDEA). The CTT sequence identity was tested using BLAST tool on [64], see the .fasta format result in S3-CTT alignment.fasta.

### Residues of drug binding sites

The paclitaxel, colchicine/nocodazole and vinca binding sites were identified based on the RSCB PDB structures 1JFF (paclitaxel is named as residue TA1, site identifier AC5) [65], 1SA0 (colchicine is named as residue CN2, site identifier AC8) [66] and 5BMV (vinblastine is named as residue VLB, site identifier AE1) [67], respectively. The residues belonging to the respective binding sites are identified there using an algorithm from [68] and are available in a respective .pdb file. For paclitaxel and colchicine, we additionally included the residues which contribute 75% binding energy between paclitaxel and tubulin based on the analysis in [69] and [70], respectively. Those results provided *β*-tubulin additional residues LEU 215, GLN 280, and LEU 361 on top of the list from RSCB PDB 1JFF structure for paclitaxel and *α*-ALA 180 and *β*-LEU 246 on top of the list from RSCB PDB 1SA0 structure for colchicine. See the complete list of residues of individual interaction and binding sites in Fig. 8.

### Molecular dynamics simulation

We carried out MD simulations of a tubulin protein in water solution using the GROMACS package v. 4.6.5 [71]. The simulated system consisted of a rectangular box (12 × 12 × 15 nm^3^), where we placed a single tubulin heterodimer, 67,087 Single Point Charge (SPC) [72] water molecules and 47 sodium ions, resulting in a typical density of 1,000 kg/m^3^. Note that, to properly describe tubulin physiological behavior, it was necessary to simulate a box of water molecules large enough to reproduce both the first hydration shells and a significant amount of bulk water. Following an energy minimization and subsequent solvent relaxation, the system was gradually heated from 50 K to 300 K using short (typically 50 ps) MD simulations. A first trajectory was propagated up to 150 ns in the NVT ensemble using an integration step of 2 fs and removing the tubulin center of mass translation but with no constraints on its related rotation. The temperature was kept constant at 300 K by the Berendsen thermostat [73] with the relaxation time equal to the simulation time step, hence virtually equivalent to the isothermal coupling [74] which provides consistent statistical mechanical behavior. All bond lengths were constrained using LINCS algorithm [75]. Long range electrostatics were computed by the Particle Mesh Ewald method [76] with 34 wave vectors in each dimension and a 4^*th*^ order cubic interpolation. The amber03 force field [77] parameters were adopted. Once obtained an exhaustive equilibrated-unexposed trajectory we evaluated possible effects induced by a high exogenous electric field ranging from 10 MV/m up to 300 MV/m (see Table 1), acting in the simulation box as explained in [78] for 30 ns in each exposure condition and applied along the z-axis of the Cartesian reference of frame. More precisely the application of the electric field takes place in continuity at the last frame of the unexposed simulation, thus allowing a direct evaluation of the characteristic response over time of the system due to the switch on of the exogenous perturbation. Taking advantages of the previous findings in [19, 56, 79] on the predictable protein polarization process, for the highest field strengths (200 MV/m and 300 MV/m) the simulation box was enlarged along the z-direction up to 30 nm, resulting in a rectangular box of 12*×*12*×*30 nm^3^.

**Table 1.** List of molecular dynamics simulations with the conditions. * the starting configuration comes from an equilibrium 150 ns trajectory (regular box, 12*×*12*×*15 nm^3^) and ** the starting configuration comes from an equilibrium 30 ns trajectory (big box, 12*×*12*×*30 nm^3^).

| Simulation number | E-field strength [MV/m] | Production time [ns] | Simulation box type |
| --- | --- | --- | --- |
| #1 | No field | 30 * | Regular |
| #2 | 10 | 30 * | Regular |
| #3 | 20 | 30 * | Regular |
| #4 | 50 | 30 * | Regular |
| #5 | 100 | 30 * | Regular |
| #6 | 200 | 10 ** | Big |
| #7 | 300 | 10 ** | Big |

### Covariance matrix method

A simple method to characterize the geometry of a complex system like a protein is to approximate its overall geometrical structure, at each MD frame, to the ellipsoid given by the eigenvectors of the Covariance Matrix as obtained by the distribution of the x, y, and z coordinates of the protein atoms (the geometrical matrix 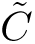) [19, 80]. Therefore, for a system defined by N-atoms the elements of the 3 × 3 geometrical matrix at each time frame are given by

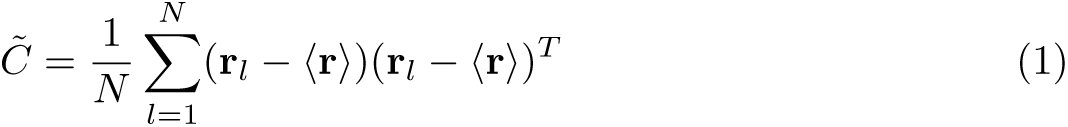

where **r**_*l*_ is the l-th atom position vector and 〈r〉 is the mean atomic position vector as obtained from averaging all the atoms position vectors. Diagonalization of the geometrical matrix provides three eigevectors (**c**_*i*_, *i* = 1, 2, 3) corresponding to the three axes of the ellipsoid best fitting the atoms distribution at that MD time frame (geometrical axes), and three corresponding eigenvalues (*λ_i_, i* = 1, 2, 3) given by the mean square fluctuations of the atomic positions along each eigenvector and providing the size of the ellipsoid along its axes.

## Supporting information

**S1-1TUB-aa content.xlsx Amino acids content in proteins** Relative content of amino acids in 1TUB tubulin versus average from proteins in UniProtKB/Swiss-Prot database

**S2-x,z dipole components Per-residue shifts of x’ and z’ dipole components**

**S3-CTT alignment.fasta Result of** *β* **tubulin CTT BLAST sequence alignment**

**S4-PDB entries.zip List of unique PDB entries**

**S5-PISCES-Charge, Dipole.zip PISCES proteins charge and dipole moments** Output from Protein Dipole Moments Server of Weizmann Institute server including charge and dipole for each pdb entry

**S6-README.txt Molecular dynamics simulation trajectories, source files and scripts** README file with with additional data and link to open-access of molecular dynamics simulation trajectories of tubulin in electric field used in this paper. Direct links are https://dx.doi.org/10.5281/zenodo.2547671 https://dx.doi.org/10.5281/zenodo.2548247 https://dx.doi.org/10.5281/zenodo.2548486

## Acknowledgements

We acknowledge Djamel Eddine Chafai, Petra Vahalová and Martin Bereta for proofreading.

## Data availability statement

See S6-README.txt for initial and configuration files as well as molecular dynamics trajectories and scripts for post-processing.

## Author contributions statement

Contribution roles according to CRediT:

http://dictionary.casrai.org/Contributor_Roles:

**Conceptualization**: Michal Cifra, Paolo Marracino, Francesca Apollonio

**Data curation**: Paolo Marracino, Michal Cifra, Francesca Apollonio, Micaela Liberti

**Formal analysis**: Paolo Marracino (lead), Michal Cifra (supporting), Jiří Průša (supporting), Daniel Havelka (supporting)

**Funding acquisition**: Lucie Kubínová, Michal Cifra

**Investigation**: Paolo Marracino (lead), Michal Cifra (supporting), Francesca Apollonio (supporting), Micaela Liberti (supporting), Jiří Průša (supporting)

**Methodology**: Paolo Marracino, Francesca Apollonio, Micaela Liberti, Michal Cifra

**Project administration**: Michal Cifra, Francesca Apollonio **Resources**: Jack Tuszynski (supporting), Ahmed T. Ayoub (supporting) **Software**: Paolo Marracino(lead), Jiří Průša (supporting)

**Supervision**: Francesca Apollonio, Michal Cifra, Micaela Liberti **Validation**: Michal Cifra, Paolo Marracino, Francesca Apollonio **Visualization**: Paolo Marracino (lead), Daniel Havelka (supporting)

**Writing – original draft**: Michal Cifra (lead), Jack Tuszynski, Paolo Marracino

**Writing – review & editing**: Michal Cifra, Francesca Apollonio, Jack Tuszynski, Micaela Liberti

